# N-Methyl-D-Aspartate receptors control *in vivo* striatal calcium and the updating of action policy

**DOI:** 10.64898/2026.07.06.736480

**Authors:** Alex A. Legaria, Mason R. Barrett, Jordyn E. Czarny, Alexxai V. Kravitz

**Affiliations:** Department of Psychiatry, Washington University in St. Louis, St. Louis MO, USA; Department of Anesthesiology, Washington University in St. Louis, St. Louis MO, USA; Department of Neuroscience, Washington University in St. Louis, St. Louis MO, USA

**Author notes:** **Corresponding Author:** Alexxai Kravitz.

**Keywords:** striatum, NMDA, calcium, reinforcement, learning, policy

## Abstract

Animals must execute learned behaviors and update them when outcomes change, yet the neural substrates controlling this phenomenon are not fully understood. Here, we show that N-Methyl-D-Aspartate Receptors (NMDARs) in the dorsomedial striatum are necessary for learning from previously rewarded actions. Moreover, blocking of striatal NMDARs almost fully abolished striatal calcium dynamics, but not action potential activity, suggesting a unique function of NMDAR-driven striatal calcium activity in updating action policy.

## Main text

Animals must select appropriate behaviors to maximize survival. To do this, they have an internal plan of actions to take in potential situations, and an ability to update these plans when the environment changes. For example, a plentiful food source may become unsafe for a mouse to forage in, requiring the mouse to adapt its behavior and forage in different places. This internal plan is known as an *action policy*^1^, which needs to be both *executed* appropriately, and also *updated* as the environment changes. The striatum, the input region of the basal ganglia, is critical for *updating* and *executing* action policies^2–4^. In particular, the dorsomedial striatum (DMS) is involved in goal-directed learning ^5^. Moreover, N-Methyl-D-Aspartate Receptors in the DMS (DMS-NMDARs) are necessary for the acquisition of new goal-directed learning^11,13–15^ and for corticostriatal plasticity^6–11^. However, whether DMS-NMDARs play a role specifically in *executing* or *updating* action policy after initial acquisition, is not clear.

We first tested the role of DMS-NMDARs in the *execution* of a previously learned action policy. To do this, we implanted mice (N=5M) with bilateral cannulas targeting the DMS and trained them on a Fixed Ratio 1 (FR1) task using a home-cage based FED3 device^12^, where nose-pokes on the correct port earned one grain pellet, while pokes on the incorrect port led to a cued 10s time out. Following training, mice were fasted for 16 hours and then infused with either saline or the non-competitive NMDAR antagonist MK-801 into the DMS (4ug/hemisphere, Figure 1A, B). Intra-DMS MK-801 did not impair execution of the previously learned FR1 task, as they performed the task with similar high accuracy (Figure 1E, p-value=0.854), and “win-stay” behavior (Figure 1F, p-value=0.706). If anything, MK-801-treated mice were slightly more engaged, earning ∼17% more pellets (Figure 1C, p-value=0.078) and performing ∼17% more pokes (Figure 1D, p-value=0.038). Overall, blocking DMS-NMDARs did not impair performance in a previously learned FR1 nose-poking task.

**Figure 1.**
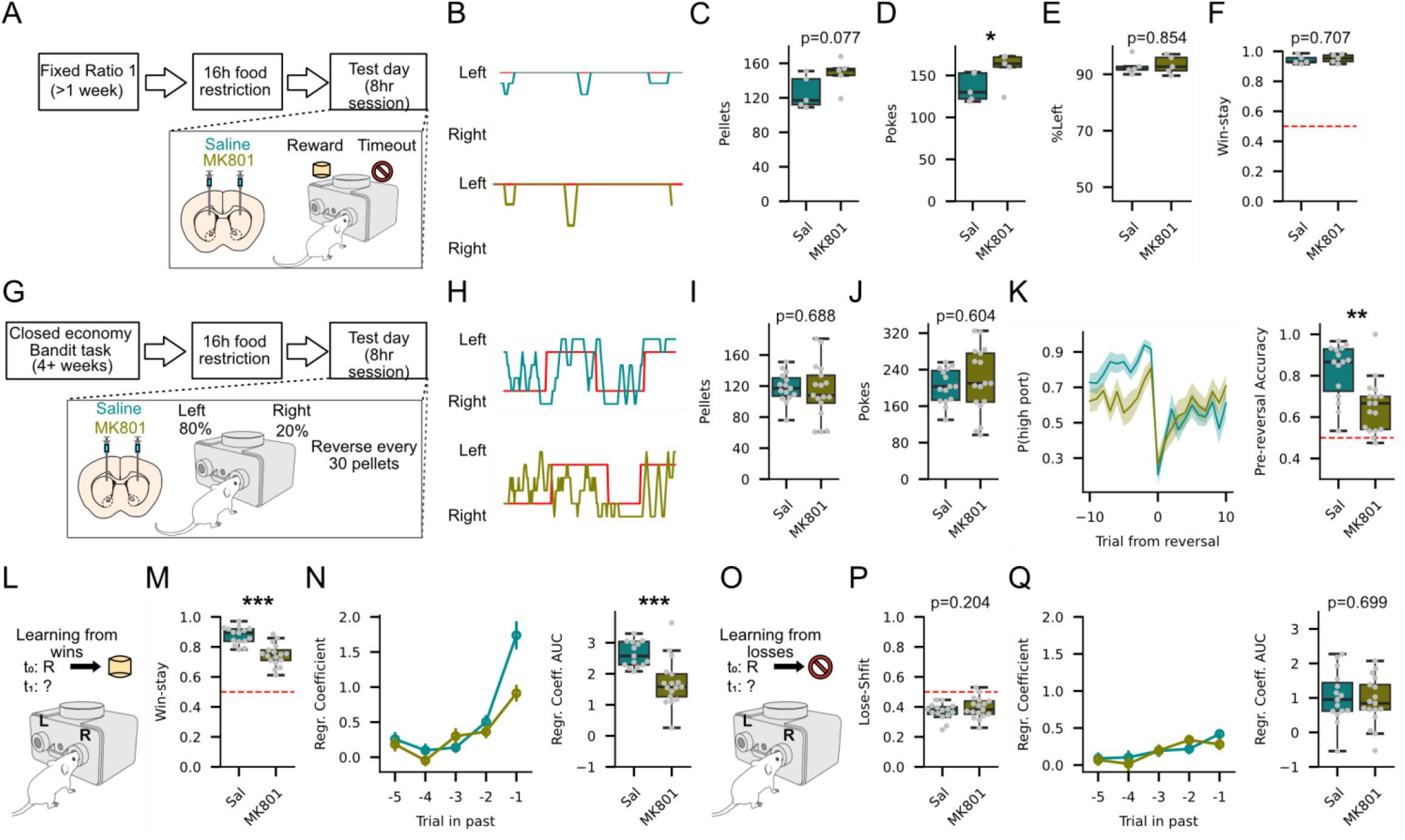
Antagonism of DMS-NMDARs disrupt update but not execution of action policy. **(A)** Experimental paradigm. Mice were trained on a Fixed Ratio 1 task. During a test session, either saline or MK-801 was infused in DMS. **(B)** Sample traces of test session after a saline infusion (top) or MK-801 infusion (bottom) in the same mouse. **(C)** Number of pellets acquired in a test session. **(D)** Number of pokes in test session. **(E)** Percentage rate of pokes on the rewarded side. **(F)** Win-stay rate. **(G)** Experimental paradigm. Mice were trained on a “two-armed” bandit task. During a test session, either saline or MK-801 was infused in DMS. **(H)** Sample traces of test session after a saline infusion (top) or MK-801 infusion (bottom) in the same mouse. **(l)** Number of pellets acquired in a test session. **(J)** Number of pokes in test session. **(K)** (Left) Peri-event histogram showing the probability of mouse poking on the “high-probability port.” Trial 0 is when the probabilities are reversed. (Right) Average accuracy (probability of poking in high-probability port) in the 10 trials prior to the reversal. **(L-N)** Assessment of leaming from rewarded trials. **(M)** Quantification of the entire test session win-stay behavior. **(N)** (Left) Regressor coefficient of logistic regression that measure influence of previously rewarded actions on choice. (Right) Sum of (left). **(O-Q)** Assessment of learning from unrewarded trials. **(O)** Quantification of lose-shift. **(Q)** Same as (I) but for unrewarded trials. Paired t-test for panels C-F. Partially-overlapping t-test for panels I-K, M,N, P, Q. * p<0.05, ** p<0.01, *** p<0.001. Full statistics including bias-corrected effect sizes in supplementary files.

Next, to test DMS-NMDARs’ role in *updating* action policy, we performed these same manipulations during a “two-armed bandit” task with FED3^13^ (21 mice 10F/11M, Figure 1G, H). Here, pokes on the high probability port had an 80% chance of delivering a pellet, and a 20% chance of delivering a 10s timeout. Pokes on the low probability port had a 20% chance of delivering a pellet and an 80% chance of delivering a timeout. The port assignments switched every 30 pellets, requiring the animals to continuously *update* their action policy to efficiently earn pellets. Intra-DMS MK-801 infusion did not significantly change the number of pellets (Figure 1I, p-value=0.688) or pokes (Figure 1J, p-value=0.604) obtained relative to saline, confirming that DMS-NMDARs are not necessary for performing the task. However, MK-801-injected mice had significantly reduced accuracy in the trials prior to port reversal (Figure 1K; ∼20% reduction, p-value=0.002), showing that mice had trouble updating their action policy after DMS-NMDAR blockade. We examined how their learning strategies changed, finding that MK-801 disrupted learning from previously rewarded actions (Figure 1L), characterized by a significant reduction (∼15%) in “win-stay” behavior (Figure 1M, p-value<0.001). Logistic regression revealed that MK-801 infusion significantly reduced (∼38%) the influence of prior rewarded trials on current choice^14–16^ (Figure 1N, p-value<0.001). However, MK-801 did not affect learning from unrewarded actions (Figure 1O), with no change in “lose-shift” behavior (Figure 1P, p-value=0.263) nor the influence of previously unrewarded trials in current choice (Figure 1Q, p-value=0.699). These effects of MK-801 were dose-dependent, with a lower dose (2ug/hemisphere) having milder effects (Extended Data Figure 1), and a systemic injection of MK-801 (1mg/kg, i.p.) showing stronger effects (Extended Data Figure 2). Together, these results demonstrate that DMS-NMDARs are critical for *updating*, but not *executing*, action policy.

We previously reported that bulk calcium striatal recordings reflect primarily dendritic calcium influx^17^. As NMDARs are a major source of dendritic calcium *ex vivo*^18,19^, we next tested whether NMDARs are implicated in modulating bulk calcium dynamics in striatal neurons. To test this, we expressed GCaMP8f in striatal neurons of the DMS of mice (N=7, 4M/3F) and used fiber photometry (with an isosbestic control signal) to record population calcium signals (Figure 2A). We injected mice with saline or the non-competitive NMDAR antagonist MK-801 (0.3mg/kg) intraperitoneally (i.p.). Following the MK-801 injection, spontaneous calcium transients were nearly completely abolished in the dorsomedial striatum (transient rate reduced by >80%, Figure 2B-D). We repeated this experiment in the Drd1-Cre (N=5, 1M/4F) and A2A-Cre (N=5M) driver lines to selectively record calcium activity in direct and indirect pathway striatal projection neurons (D1-SPNs and D2-SPNs), again observing near-complete cessation (>90% reduction) of spontaneous calcium transients in both populations (Figure 2B-D, p-value<0.001 for both D1-SPNs and D2-SPNs). We conclude that NMDARs are necessary for spontaneous calcium dynamics in the DMS.

**Figure 2.**
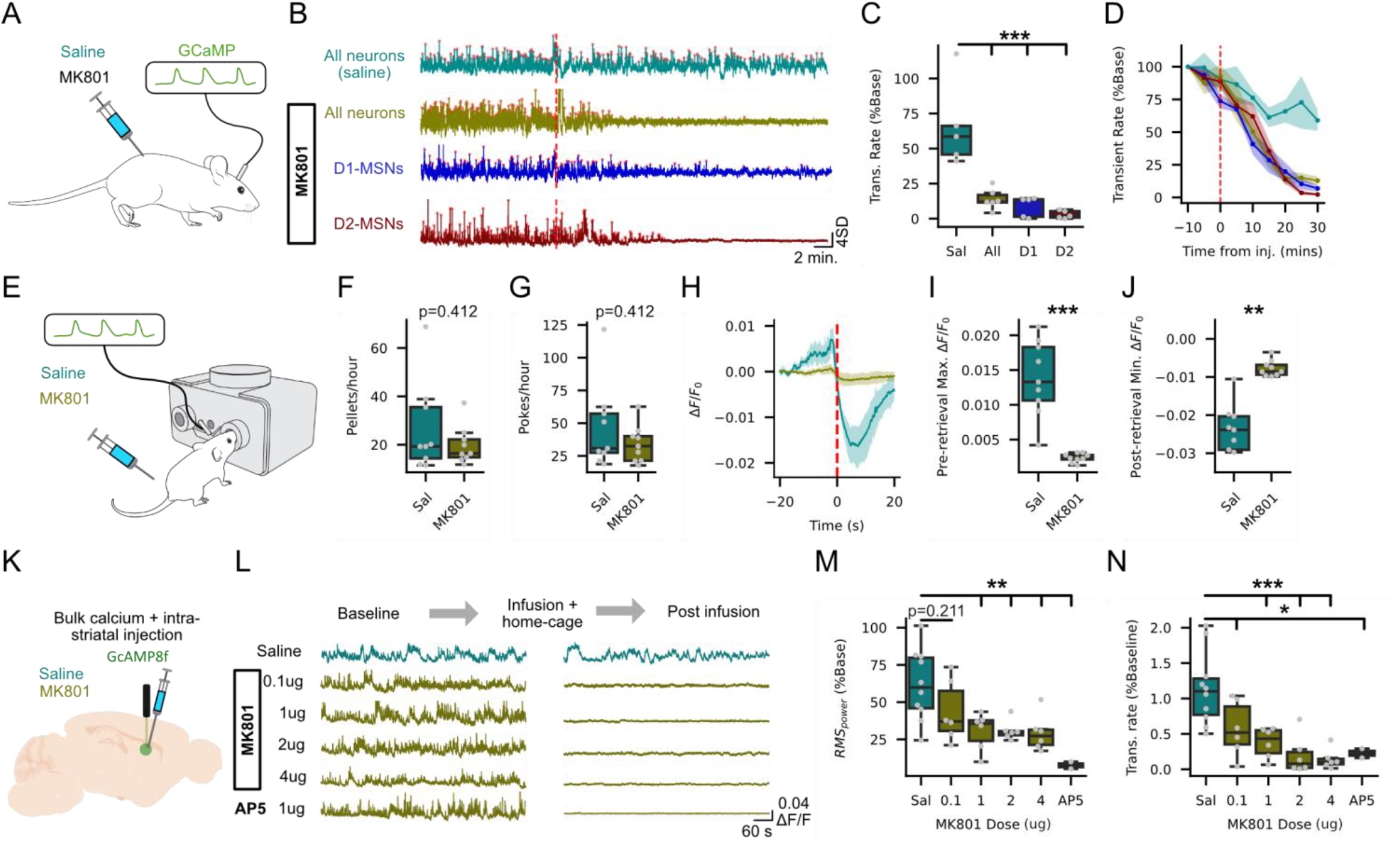
Striatal calcium dynamics are NMDAR-dependent and are not necessary for action policy execution. **(A)** Bulk striatal calcium activity was recorded before and after i.p. injection of the non-competitive antagonist MK-801. **(B)** Example calcium activity traces after saline or MK-801 injection in all striatal populations, D1-MSNs, or D2-MSNs. Red dotted line indicates time of injection. Red dots indicate identified peaks in calcium activity (“transients”). **(C)** Quantification of the transient rate for the last 10 minutes (20 to 30 minutes post-injection). **(D)** Transient rate time course in 5 minute bins around saline or MK-801 injection, normalized to the first 5 minutes of the recording. **(E)** Mice were trained on an operant task, where they nose-poked on a feeding device to obtain a food reward. Saline or MK-801 was injected i.p. while bulk calcium activity in the striatum was recorded. **(F)** Calcium activity around pellet retrieval. **(G)** Quantification of pre-retrieval response. **(H)** Quantification of post-retrieval inhibition. **(l)** Pellet rate. **(J)** Poke rate. **(K)** Bulk calcium recordings were coupled with local infusions of saline, MK-801, or APV **(L)** Example traces of calcium activity before infusion (baseline) and after infusion (Post infusion) of saline, different doses of MK-801, or APV. All the traces are from the same mouse in different sessions. **(M)** Quantification of transient rate post infusion, normalized to the baseline period transient rate. One-way ANOVA (F-stat=16.9, p-value<0.001) with post-hoc Dunnett test for panel C. Paired t-test for panels F,G,I,J. Linear mixed model with post-hoc Dunnett test for panels M.N (Linear mixed model p-values<0.001 for all groups in both M and N). p<0.05, ** p<0.01, *** p<0.001. Full statistics including bias-corrected effect sizes in supplementary files.

We next tested if NMDARs are also necessary for behaviorally evoked calcium signaling in the FR1 task. After training on the FR1 task, we injected D1-SPNs (N=5, 3M/2F) or D2-SPNs (N=5M) mice with saline or MK-801 (0.3mg/kg i.p.) and recorded GCaMP8f signals while they executed the FR1 task (Figure 2E). In saline-injected mice, calcium activity ramped up over ∼15 seconds prior to the retrieval of the pellet and was inhibited below baseline while the mouse consumed the pellet, consistent with previous reports^20^ (Figure 2H). However, both the increase in calcium prior to pellet retrieval and the reduction post-retrieval were nearly completely abolished in MK-801 treated mice compared to saline (Figure 2H-J; ∼86% reduction in pre-retrieval response, p-value<0.001. ∼68% reduction in post-retrieval inhibition, p-value=0.001). Similar results were obtained in both D1-SPNs and D2-SPNs, and around different behavioral motifs within trials (Extended Data Figure 3). These results show that both spontaneous and behaviorally evoked calcium dynamics in the DMS require NMDARs.

An important caveat to the above calcium recordings was the systemic administration of MK-801, which may reduce excitatory drive across the entire brain. To test the effect of local striatal NMDAR blockade, we performed microinjections of MK-801 into the DMS while simultaneously recording calcium activity. Our injection volume was 1uL, which has an estimated pharmacological spread of ∼1mm ^21,22^ (Figure 2K, N=12, 4F/ 8M). 1ug, 2ug, and 4ug of MK-801, and the competitive NMDA antagonist AP5 microinjection strongly reduced the power of calcium dynamics in the DMS (Figure 2L,M; reductions relative to saline: 0.1ug ≈30%, p-value=0.210; 1ug≈50%, p-value=0.007; 2ug≈50%,p-value=0.008; 4ug≈52% p-value=0.005; AP5≈88%, p-value=0.002). This effect was dose-dependent (Z-value=-5.340, P-value<0.001). Similarly, all doses of MK-801 and AP5 significantly reduced the frequency of spontaneous transients (Figure 2N, reductions relative to saline: 0.1ug≈50%, p-value=0.021 1ug≈67%, p-value=0.001; 2ug≈83%,p-value<0.001; 4ug≈88%, p-value<0.005; AP5≈81%, p-value=0.012). Similar to signal power, the MK-801 effect was dose-dependent (Z-value=-7.032, P-value<0.001), with the 2ug and 4ug doses reducing the transient rate to similar levels as with i.p. injection. We also performed a spectral analysis of the entire photometry recording following the infusion, finding a broad-band reduction in power across all frequencies (Extended Data Figure 4). We conclude that striatal NMDARs control striatal calcium dynamics, and that antagonism of these receptors strongly abolishes striatal calcium dynamics.

Finally, we tested the possibility that the behavioral and physiological effects of MK-801 may be driven by a disruption in action potential activity, since NMDARs are also important for bursting activity of striatal neurons^23^ and backpropagating (BP) action potentials have been proposed as a source of the non-somatic changes observed in bulk calcium recordings *in vivo*^24^. We performed simultaneous *in vivo* electrophysiology and bulk calcium recordings^17^ while exposing mice to MK-801 (Figure 3A, N=6, 2F/4M). Again, MK-801 administration (0.3mg/kg, i.p.) nearly completely (>90%) eliminated calcium transients (Figure 3B,C). MK-801 injection also reduced the frequency of firing bursts (∼53%), but to a lesser degree than calcium transients (Figure 3C, p-value=0.008). This reduction in firing bursts was not observed after a saline injection, although calcium transients were partially reduced, possibly due to photo bleaching. Nevertheless, the reduction in calcium transients was significantly greater after MK-801 injection compared to saline injection (Extended Data Figure 5).

**Figure 3.**
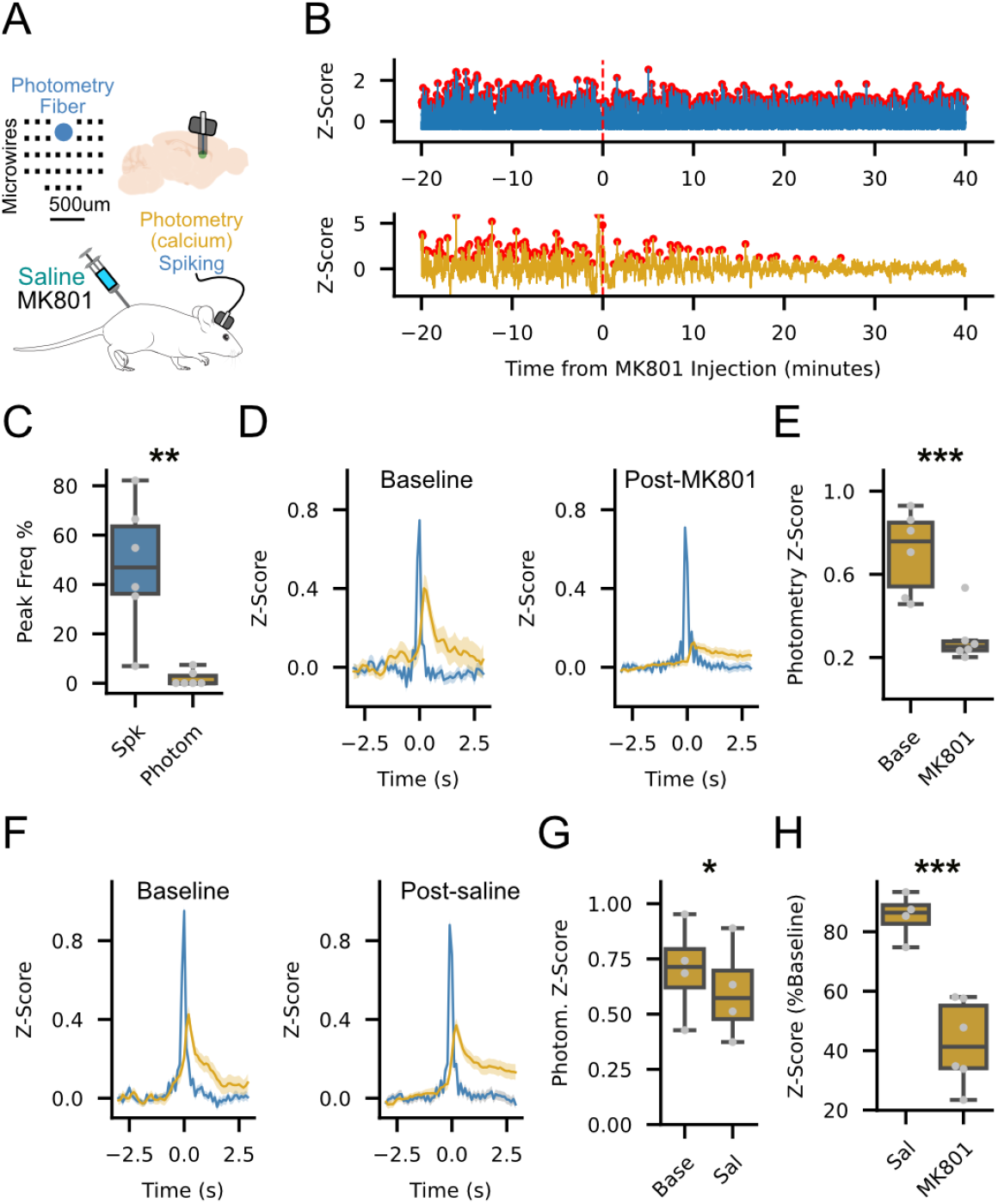
NMDAR antagonism dissociates striatal calcium dynamics from action potential activity. **(A)** Simultaneous *in vivo* electrophysiology and calcium recordings before and after systemic IMK-801 injection. **(B)** Example trace of simultaneous average firing rate (top) and calcium activity (bottom) before and after MK-801 injection. **(C)** Frequency of rapid increase in firing rate (Bursts) or calcium activity (Transients), normalized to the baseline period. **(D)** Firing bursts of similar magnitude during the baseline (left) and post-MK801 (right) period. Blue trace displays firing rate activity during the burst, and the gold trace displays the concurrent calcium activity. **(E)** Quantification of imaximum photometry response around bursts. Each dot represents the average maximum response per mouse. **(F)** Same as D for saline iinjection. **(G)** Same as (E) for the saline injection. **(H)** Comparison of calcium active after saline (from (G)) or MK801 response (from (E)) around bursts. Paired t-tests for panels C,E,G. Independent t-test for panel H. *p<0.05, ** p<0.01, *** p<0.001. Full statistics including bias-corrected effect sizes in supplementary files.

To test if the reduction in calcium dynamics reflected the reduction in bursting, we analyzed calcium rises around firing bursts. Before the MK-801 administration, action potential bursts were associated with rises in population calcium (Figure 3D). However, when we identified bursts of matching magnitudes after MK-801 infusion, these bursts were associated with a significantly smaller rise in calcium activity (Figure 3E, ∼60% reduction relative to the baseline period, p=0.002). As a control, we repeated these experiments with a saline injection instead of MK-801. Calcium rises around bursts were maintained after a saline injection compared to baseline, with only a small, yet significant decrease (Figure 3F, G, ∼14% reduction, p-value=0.035), again possibly due to photo bleaching. Nevertheless, the reductions in calcium around bursts were significantly greater after MK-801 compared to saline injection (Figure 3H, p-value<0.001). As such, calcium diminishment by NMDAR antagonism persisted even when bursting activity is preserved, suggesting that calcium activity can be dissociated from action potential activity.

Here, we found that DMS-NMDARs are necessary for *updating* action policy but are dispensable for its *execution*. Moreover, we found that NMDARs are required for striatal calcium dynamics, which can be dissociated from action potential activity. These results help interpret striatal fiber photometry in a new light. While commonly interpreted as a proxy for action potential activity, or generic “neural activity,” we instead show that these dynamics reflect NMDAR-dependent dendritic calcium influx, which can be regulated independently from action potential firing.

Integrating our two major observations, striatal NMDAR-mediated calcium dynamics are well-posed to be a physiological substrate linking plasticity events to action policy update, possibly functioning as an eligibility trace through a three-factor learning rule^25^.

Our study has limitations. While we show that antagonism of NMDARs is necessary for both bulk striatal calcium dynamics and action policy *updating*, we do not show a direct link between calcium levels and policy *updating* on a trial-by-trial basis. In fact, during the FR1 behavioral task, calcium levels were highest near the pellet retrieval (Extended Data Figure 3F), and relatively low when the outcome tone occurred (Extended Data Figure 3D), which is when dopamine error signals are predicted to occur. This suggests a more complex relationship between learning (and/or plasticity) than the product of calcium and dopamine levels at the time of phasic dopamine release, or that heterogeneity of local calcium levels cannot be detected by fiber photometry due to its limited spatial resolution.

However, recent studies have shown evidence of information about recent action-outcome associations present in calcium activity of SPNs, which can causally direct future choice^26^. Moreover, more experiments are necessary to fully dissect the *in vivo* relationship between NMDAR-driven calcium activity and action potential activity. Finally, we attempted to fit reinforcement learning models to our data, but our freely moving behavioral paradigm only provided ∼100-200 trials per mouse, which did not allow for consistent convergence of fitted reinforcement learning models. This issue would need to be examined with other behavioral tasks in the future that produce more trials. Despite these limitations, our study advances our understanding of the *in vivo* role of striatal NMDARs in learning and calcium signaling and further cements the view of the striatum as a hub for reinforcement learning in the brain.

## Methods

All experimental procedures were approved by the Washington University Animal Care and Use Committee and the Northwestern University Animal Care and Use Committee.

### Subjects

The animals used in this study were 40 wild-type C57BL6 mice (23 males, 17 females), 9 Drd1-Cre mice (GENSAT line EY217, 5 females, 4 males), 8 A2a-Cre mice (GENSAT line KG139, 8 males). Cre mice were obtained from the GENSAT project (The Gene Expression Nervous System Atlas (GENSAT) Project, NINDS Contracts N01NS02331 & HHSN271200723701C to The Rockefeller University (New York, NY)). Animals were housed in the Washington University in St. Louis animal facilities in standard vivarium cages with ad libitum food and water and a non-reversed 12-h dark/light cycle.

### Surgery

#### Viral transduction

Anesthesia was induced with 3-5% isoflurane and maintained at 0.5–1.5% isoflurane during stereotactic surgery. Ear bars and a mouth holder were used to keep the mouse head in place while the skin was shaved and disinfected with a povidone/iodine solution. The skull was exposed and 1-mm-diameter craniotomy was made with a microdrill mounted to the stereotaxic manipulator. Injections were performed with a glass pipette mounted in a Nanoject 3 infusion system (Drummond Scientific). Then 350 nL of AAV1-Syn-GCaMP8f or AAV2/9-hSyn-FLEX-GCaMP8f (1.2 × 10^12^ GC ml^−1^, Addgene plasmid #162379 and #162376, respectively) virus was infused over 6 min into the dorsal striatum (AP = +0.5 mm, ML = +1.5 mm, DV = −2.8 mm). The injector was left in place for 5 min before removal.

#### Implantation of fiber photometry cannula

For fiber photometry experiments that only required an optical fiber, after viral infusion a fiber photometry cannula (Neurophotometrics or RWD, 5mm fiber length from ferrule, 200 um diameter, 0.5 numerical aperture) was lowered to 0.1 mm above the viral infusion coordinates. A layer of Metabond (C&B Metabond, Parkell) was placed on the skull and enough acrylic dental cement (Lang dental) was added to secure the fiber in place. Once the cement had fully cured, animals received a subcutaneous injection of meloxicam (10 mg kg^−1^) and placed back in their home-cage on a pre-heated pad at 37 °C. Mice recovered for at least 2 weeks to allow for viral expression before recording.

#### Implantation of electrode array and optical fiber cannula bundle

For combined electrophysiology and fiber photometry experiments, the fiber photometry cannulas (Neurophotometrics or RWD, 5mm fiber length from ferrule, 200 um diameter, 0.5 numerical aperture) were mounted in a custom electrode array with 32 Teflon-coated tungsten microwires (35 µm diameter; Innovative Neurophysiology) that positioned the wires in a semicircle surrounding a central gap where the photometry fiber was mounted. After viral infusion, a larger craniotomy was performed to allow implantation of the array. The combined photometry/electrical recording device was implanted into the right DMS (AP = +0.5 mm, ML = +1.5 mm, DV = −2.7 mm). The device was secured to the skull with a thin layer of adhesive dental cement (C&B Metabond, Parkell) followed by a larger layer of acrylic dental cement (Lang Dental). Once the cement had fully cured, animals received a subcutaneous injection of meloxicam (10 mg kg^−1^) and placed back in their home-cage on a pre-heated pad at 37 °C. Mice recovered for at least 2 weeks to allow for viral expression before recording.

#### Implantation of infusion cannula and optical fiber cannula bundle

For intra-striatal infusion near the fiber photometry field of view, a guide cannula (26G, ∼3.5mm length), was mounted on a fiber photometry cannula (Neurophotometrics or RWD, 5mm fiber length from ferrule, 200 um diameter, 0.5 numerical aperture) using UV light-cured epoxy. Particular care was taken to verify that the end of the guide cannula was as close as possible to the end of the optical fiber cannula. A silver wire was inserted into the guide cannula to prevent clogging. After viral infusion, the infusion cannula/optical fiber bundle was lowered to 0.1 mm above the viral infusion coordinates. A layer of Metabond (C&B Metabond, Parkell) was placed on the skull and enough acrylic dental cement (Lang dental) was added to secure the fiber in place. Once the cement had fully cured, animals received a subcutaneous injection of meloxicam (10 mg kg^−1^) and placed back in their home-cage on a pre-heated pad at 37 °C. Mice recovered for at least 2 weeks to allow for viral expression before recording.

#### Implantation of bilateral cannulas

Anesthesia was induced with 3-5% isoflurane and maintained at 0.5–1.5% isoflurane during stereotactic surgery. Ear bars and a mouth holder were used to keep the mouse head in place while the skin was shaved and disinfected with a povidone/iodine solution. The skull was exposed and 0.5-mm-diameter craniotomies were made in either hemisphere with a microdrill mounted to the stereotaxic manipulator at coordinates AP: +0.5, ML: ±1.5. Then, the bilateral guide cannula (26G, 3mm cannula length, 3mm center-to-center, 5.5 mm total length including cannula base) was lowered to -2.8mm below the skull through the craniotomies. A layer of Metabond (C&B Metabond, Parkell) was placed on the skull and enough acrylic dental cement (Lang dental) was added to secure cannula in place. Once the cement had fully cured, animals received a subcutaneous injection of meloxicam (10 mg kg^−1^) and placed back in their home-cage on a pre-heated pad at 37 °C. Mice recovered for at least 2 weeks to allow for viral expression before recording.

### Pharmacology

#### Intraperitoneal injections

Intraperitoneal injections of MK-801 were done at a dose of 0.3mg/kg for Figure 2(A-J), Figure 3, and Extended Data Figure 3 and 5, and of 1mg/kg for Extended Data Figure 2, from 0.03mg/mL MK-801 or 0.1mg/mL MK-801 solution, respectively, with a volume of 10uL per gram of body weight. Similarly, saline injection volume was calculated at 10uL per gram of body weight. After infusion, mice were allowed to recover for 10 minutes and placed either back into the recording chamber (Figure 2A-J, Extended Data Figure 3) or the FED3 device was placed back in the home-cage (Extended Data Figure 2). For Figure 3, mice were placed back in the recording chamber immediately after injection.

#### Intra-striatal infusions

Intra-striatal infusions were performed using a syringe pump (Legato 101 syringe pump, order code: 78810). Hamilton syringes (5uL) were mounted to the pump and connected to an injection cannula (33G, 3.00mm length) via BTPE tubing (Instech BTPE-50PE-50 tubing, fits 22ga, .023x.038in.). The flow rate of all intra-striatal infusions was 0.2uL/min, and the total volume was 1uL. First, the Hamilton syringes were loaded with saline or MK-801. Then, the mice were scruffed and the dummy probe (or silver wire) was removed. The injection cannula was then inserted into the guide cannula, and the mice were placed back in their open (no lid) home cage. Infusion was then started, taking care that there was no tangling of the tubing. After infusion, mice were allowed to recover for 10-30 minutes and placed either back into the recording chamber (Figure 2K-N and Extended Data Figure 4) or the FED3 device was placed back to the home-cage (Figure 1, Extended Data Figure 1).

### Neural recordings

#### Fiber photometry recordings

Fiber photometry recordings were performed in a FP3001 fiber photometry system (Neurophotometrics). Briefly, this system utilizes a 470-nm blue-light LED, which was left on continuously at 40– 100 μW to measure calcium-dependent GCaMP signals and a 405-nm UV light was used to measure calcium-independent GCaMP emissions. The light path includes a dichroic mirror to pass emitted fluorescence to a complementary metal-oxide semiconductor camera (FLIR BlackFly). Fluorescence signals from the camera were collected and processed with Bonsai (https://bonsai-rx.org/docs).

#### Electrophysiological recordings

Neurophysiological signals were recorded by a multichannel neurophysiology system (Plexon Omniplex, Plexon Inc.). Spike channels were acquired at 40 kHz and bandpass filtered from 150 Hz to 3 kHz before spike sorting. Recordings were performed in a 12” × 12” white acrylic box and lasted between 1 and 2 hours. Video and tracking data was also recorded in real time with the Plexon Cineplex system.

#### Simultaneous Fiber photometry and electrophysiological recordings

Fiber photometry and electrophysiology data was collected as previously described. Additionally, fiber photometry data was transmitted through a DAC as a voltage signal to the Plexon Omniplex for simultaneous digitizing with the electrophysiological data.

### Behavior

#### Open Field

For experiments where only spontaneous neural activity was recorded (Figure 2A-D, Figure 3, Extended Data Figure 4,5), mice were placed in an open field chamber and allowed to freely move while neural activity was recorded, in 1–2-hour sessions.

#### Fixed Ratio 1 task

For training mice on a fixed ratio 1 task, a Feeding Experimentation Device version 3 (FED3) was placed in the home cage. The FED3 dispensed 20mg grain pellets (Bio-serv F0163 “Dustless Precision Pellets^®^ Rodent, Grain-Based), which were the only food source in the mouse’s cage. To obtain pellets, mice were required to nose-poke on the left nose-port (left poke). One second after a left poke, a 4kHz tone played for 200ms indicated a successful trial. If the mouse poked on the right nose-port (right poke), a white noise tone was played for 10 seconds, which indicated an unsuccessful trial. Two seconds after a successful trial, a pellet was delivered.

Mice were trained on this task for at least 3 days (range 3 days - >2 weeks). All mice successfully obtained all their food through the FED3 (at least >100 pellets per day), with the vast majority of pokes occurring on the rewarded side.

After mice had been trained, the night prior to the behavioral session, the FED3 was removed from the home cage, inducing food restriction (between 16-24 hours). Prior to the behavioral session, mice were treated with saline or MK-801, following the aforementioned procedure, with intra-striatal infusions for experiments on Figure 1A-F, and i.p. injections for experiments on Extended Data Figure 2A-F. Ten minutes after, mice were placed in their home cage (Figure 1A-F and Extended Data Figure 2 A-F) or recording chamber (Figure 2E-J) and were allowed access to the FED3.

#### Two-armed bandit task

For training mice on the two-armed bandit task, a FED3 was placed in the home cage. The FED3 dispensed the 20mg grain pellets which were the only food source in the mouse’s cage. First, mice were trained on a deterministic reversal task. In this task, one of the nose-ports on one side (e.g. Left) of the device was always associated with a “rewarded” tone (4kHz for 200ms) and the delivery of a food pellet, while the other nose-port was always associated with an “unrewarded” tone (white noise for 10 seconds) and a 10 second timeout, where mice could not receive a reward upon any nose-poke. Importantly, this time out co-occurred with white noise and reset if the mouse poked on any nose-port. The pokes associated with reward and timeout reversed every time the mice received 30 pellets. Mice were considered to have “learned” the task, if the average accuracy (percent of pokes done in the rewarded poke) over the last 3 days of training reached at least 70%, with a minimum time of training of 1 week, and a maximum of 3 weeks. Mice that reached this metric, were moved onto the bandit task, where instead of one side being always associated with reward and the other always to a time out, these associations were probabilistic. The nose-port on one side (e.g. Left) had an 80% probability of leading to a “rewarded” tone and food delivery, and a 20% probability of leading to an “unrewarded” tone and a timeout. These probabilities reversed every time the mouse obtained 30 pellets. Mice were considered to have “learned” the bandit-task, if the average accuracy (percent of pokes done in the high-probability poke) over the last 3 days of training reached at least 60%. Mice were trained on the bandit-task for at least 2 weeks before moving onto the test session.

The night prior to the test session, the FED3 was removed from the home cage, inducing food restriction (between 16-24 hours). Prior to the behavioral session, mice were treated with saline or MK-801, following the aforementioned procedure (see pharmacology section) with intra-striatal infusions for experiments in Figure 1G-Q, and Extended Data Figure 1, and i.p. injections for experiments in Extended Data Figure 2G-Q. Ten minutes after delivery of saline or MK801, mice were placed in their home cage and were allowed access to the FED3 device. Data spanning eight hours starting from when the mouse poked for the first time after treatment were analyzed.

### Data analysis

All analyses (except spike sorting) were performed using custom Python pipelines. All the analysis code is available at: https://github.com/AlexLM96/Legaria_etal_2025

#### Photometry signal preprocessing

Noise in the photometry signal was approximated by fitting the isosbestic (calcium-independent) signal to the calcium-dependent signal using a linear regression. The ΔF/F0 was calculated using the following equation:

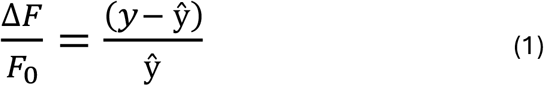

where *y*is the measured calcium-dependent signal and ŷ is the predicted signal from the linear regression. Then, a Butterworth bandpass filter was applied to the resulting signal, with low and high critical frequencies of 0.005Hz and 10 Hz, respectively, to correct for noise and photo bleaching. Finally, the signal was downsampled to 10Hz.

The following equation was used to calculate Z-Score:

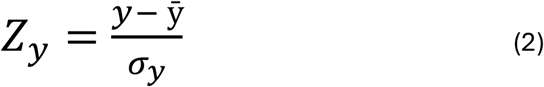

Where *y* is the pre-processed GCaMP signal.

#### Population spiking preprocessing

Single units and multi-units were manually sorted using Plexon Offline Sorter. To calculate population spiking, first firing rate of each unit was calculated in 100ms bins. The resulting signal was convolved with a 300ms boxcar window. Each unit was then z-scored to itself, and all the units were averaged.

#### Photometry transient and spiking burst detection

Photometry transients were detected using two methods. For Figure 2N, photometry transients were detected through wavelet convolution, using the function find_peaks_cwt() from the python scipy.signal module. The exact parameters used can be found in the Python pipelines (https://github.com/AlexLM96/Legaria_etal_2025).

For Figure 2B-D and Figure 3 B-C, photometry transients were detected using the scipy.find_peaks() function, setting a minimum prominence of 1.5 for calcium transients and 1 for spiking bursts. This prominence-based method to detect calcium transients was used to ensure consistency in peak identification, since wavelet convolution did not lead to reliable identification of spiking bursts.

#### Peri-event histograms

For all peri-event histograms, each trial was normalized to a baseline window by subtracting the mean of that trial baseline. For instance, in Figure 2H the baseline window spanned (-20 to -15 seconds). For each trial, the mean of the -20 to -15 seconds was calculated, and this mean was subtracted to the entire trial. After normalization, all trials were averaged.

#### Power analysis

To calculate the root-mean-squared average power of the photometry signals (Figure 2M), the following equation was used:

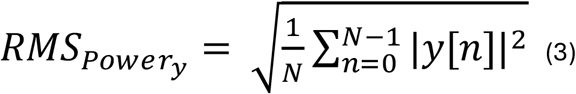

where y is the processed GCaMP signal. To calculate the frequency-based power (Extended Data Figure 4), Welch’s method was used (from python’s scipy.signal.welch). The specific parameters can be found in the python pipeline.

### Analysis of behavioral data

#### Accuracy peri-event histogram

For every probability reversal, the ten trials prior and posterior to the reversal were used. If the mouse poked on the side associated with a high probability of reward, then a value of 1 was assigned to that trial, and a value of 0 otherwise, resulting in an 11-element array of 1s and 0s for each trial. All the trials for each mouse were averaged.

#### Win-stay and lose shift

For calculating the win-stay metric, for every trial when the mouse received a reward (*C*(*i*),*when R* = 1), if during the next trial the choice (*C*(*i*+1)) was repeated (*C*(*i*+1)= *C*(*i*)), then that trial was assigned a value of one (*WS*(*i*)= 1), and 0 otherwise (*WS* (*i*)=0). The win-stay metric was the average of the value of all the trials 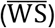. For lose-shift, the same procedure was followed for every unrewarded trial (*C*(*i*),*when R* =0), where the trial was assigned a value of one if the choice was switched after the unrewarded trial, otherwise a value of zero was assigned (*LS*(*i*)= 1, *if C*(*i*+1)= *C*(*i*),*otherwise LS*(*i*)=0 . The win-stay metric was the average of the value of all the trials 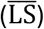.

#### Logistic Regression

Logistic regression was applied to predict each mouse’s choice as a function of previous choice and outcome history. The model uses the rewarded choice history *R*(*i* −*j*)(equation 4) and the unrewarded choice history *N*(*i* −*j*)(equation 5) to predict choice log 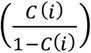, where *C*(*i*)= 1 for a left choice and 0 for a right choice.

*R* = 1 for a rewarded choice and 0 for an unrewarded choice, and *N* = 1 for an unrewarded choice and 0 for a rewarded choice. A positive regression coefficient suggests the tendency to repeat previous action, while a negative regression coefficient indicates the likelihood of switching to the other nose-port. Logistic regression was fit using maximum likelihood estimation in Python’s Stats models.

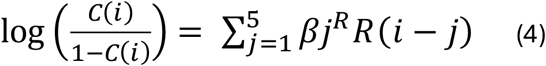

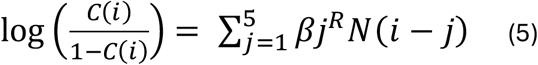

### Histology

At the end of the experiments, we performed histological verification of implant placements. Animals were anesthetized with isoflurane and decapitated, and their brains were quickly removed and placed in 10% formalin solution, in which they were incubated overnight. The brains were then moved to 30% sucrose solution until sectioning. Coronal slices containing the striatum were prepared using a freezing microtome (Leica SM2010R). Slices were mounted on microscope slides with a mounting media and imaged with an epifluorescence microscope (Zeiss). For in vivo electrophysiology, electrode placement was assessed via observation of implant tracts or electric lesions that were made under anesthesia before decapitation (performed with a 5 -s long pulse of 10 mA; Ugo Basile Lesion Making Device).

### Rigor and Reproducibility

No statistical methods were used to pre-determine sample sizes, but our sample sizes are similar to those reported in previous publications.

Confirmation of viral expression and optic fiber/electrode implantation in each mouse was done through histology (see Histology section) and functionally (spontaneous calcium transients clearly observable by eye). For behavioral experiments, only mice that reached the pre-established performance metrics during training (see two-armed bandit task subsection) were used for further experiments. A preliminary cohort of mice was used to prepare analysis pipelines, and subsequent experiments were run through this pipeline. Experimenters were not blinded to the identity of compounds (saline or MK-801) injected but all files were run indiscriminately through the same analysis pipeline. No data were excluded.

## Data Availability

The datasets generated during and/or analyzed during the current study are available in the Open Science Framework (OSF) repository, at https://osf.io/stk2r/files.

## Code Availability

All custom code generated to analyze the datasets used in the current study is available on GitHub at: https://github.com/AlexLM96/Legaria_etal_2025.

## Acknowledgements

This work was supported by the NIH R01DK136810 (AVK), DK138131 (AVK), the Washington University Diabetes Research Center Pilot Award Program (AVK), the Washington University Nutrition Obesity Research Center Pilot Award Program (AVK), and the Taylor Family Institute for Innovative Psychiatric Research, Washington University School of Medicine.

## Notes

https://github.com/AlexLM96/Legaria_etal_2025.

### Competing Interest Statement

The authors have declared no competing interest.

