## Extended Data Figure for "N-Methyl-D-Aspartate receptors control *in vivo* striatal calcium and the updating of action policy"

### Extended Data Figure 1

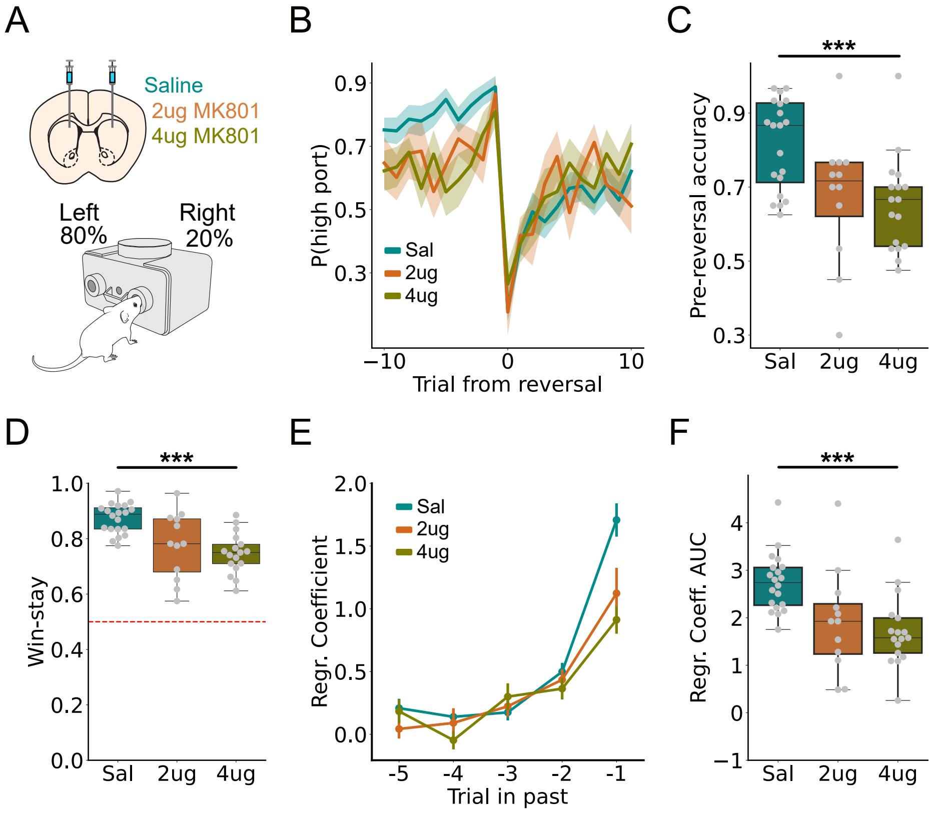


**Extended Data Figure 1. Learning effects of DMS-NMDAR antagonism is dose-dependent. (A)** Experimental paradigm. Same as Figure 1G-Q. Either saline or one of two doses of MK-801 were infused into the striatum. **(B)** (Left) Peri-event histogram showing the probability of mouse poking on the "high-probability port." Trial 0 is when the probabilities are reversed. **(C)** Average accuracy (probability of poking in high-probability port) in the 10 trials prior to the reversal. **(D)** Quantification of the entire test session win-stay behavior. **(E)** Regressor coefficients of logistic regression that measure influence of previously rewarded actions on choice. **(F)** Sum of coefficients in (E).

### Extended Data Figure 2

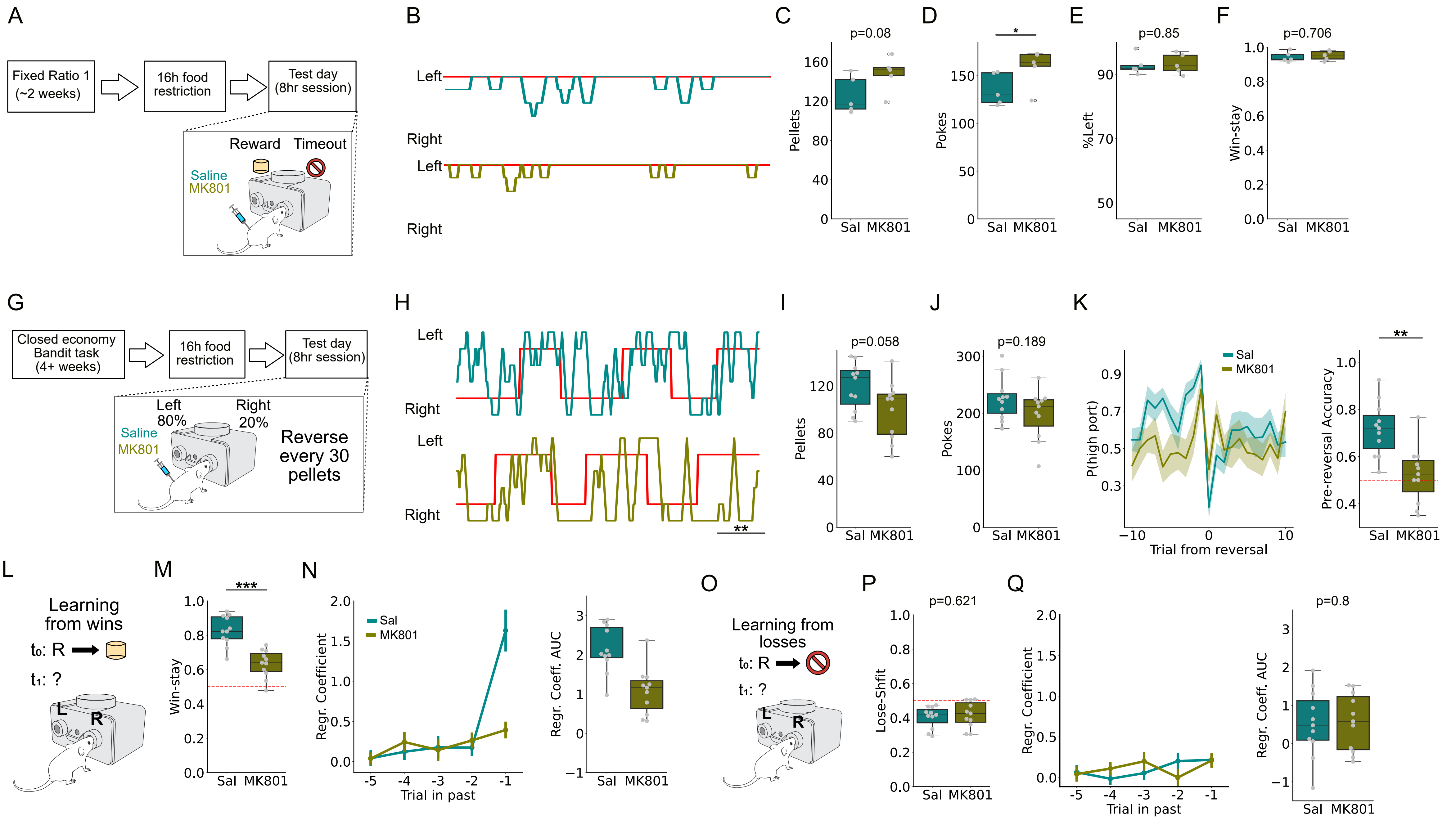


**Extended Data Figure 2. Systemic antagonism of NMDARs disrupt update but not execution of action policy. (A)** Fixed ratio 1 experimental paradigm. **(B)** Sample traces of test session after a saline (top) or MK-801 injection (bottom) in the same mouse. **(C)** Number of pellets acquired in a test session. **(D)** Number of pokes in test session. **(E)** Percentage rate of pokes on the rewarded side. **(F)** Win-stay rate. **(G)** Two-armed band task experimental paradigm. (H) Sample traces of test session after a saline infusion (top) or MK-801 infusion (bottom) in the same mouse. **(I)** Number of pellets acquired in a test session. **(J)** Number of pokes in test session. **(K)** (Left) Peri-event histogram showing the probability of mouse poking on the "high-probability port." Trial 0 is when the probabilities are reversed. (Right) Average accuracy (probability of poking in high-probability port) in the 10 trials prior to the reversal. **(L-N)** Assessment of learning from rewarded trials. **(M)** Quantification of the entire test session win-stay behavior. **(N)** (Left) Regressor coefficient of logistic regression that measure influence of previously rewarded actions on choice. (Right) Sum of (left). **(O-Q)** Assessment of learning from unrewarded trials. **(P)** Quantification of lose-shift. **(Q)** Same as (N) but for unrewarded trials.

### Extended Data Figure 3

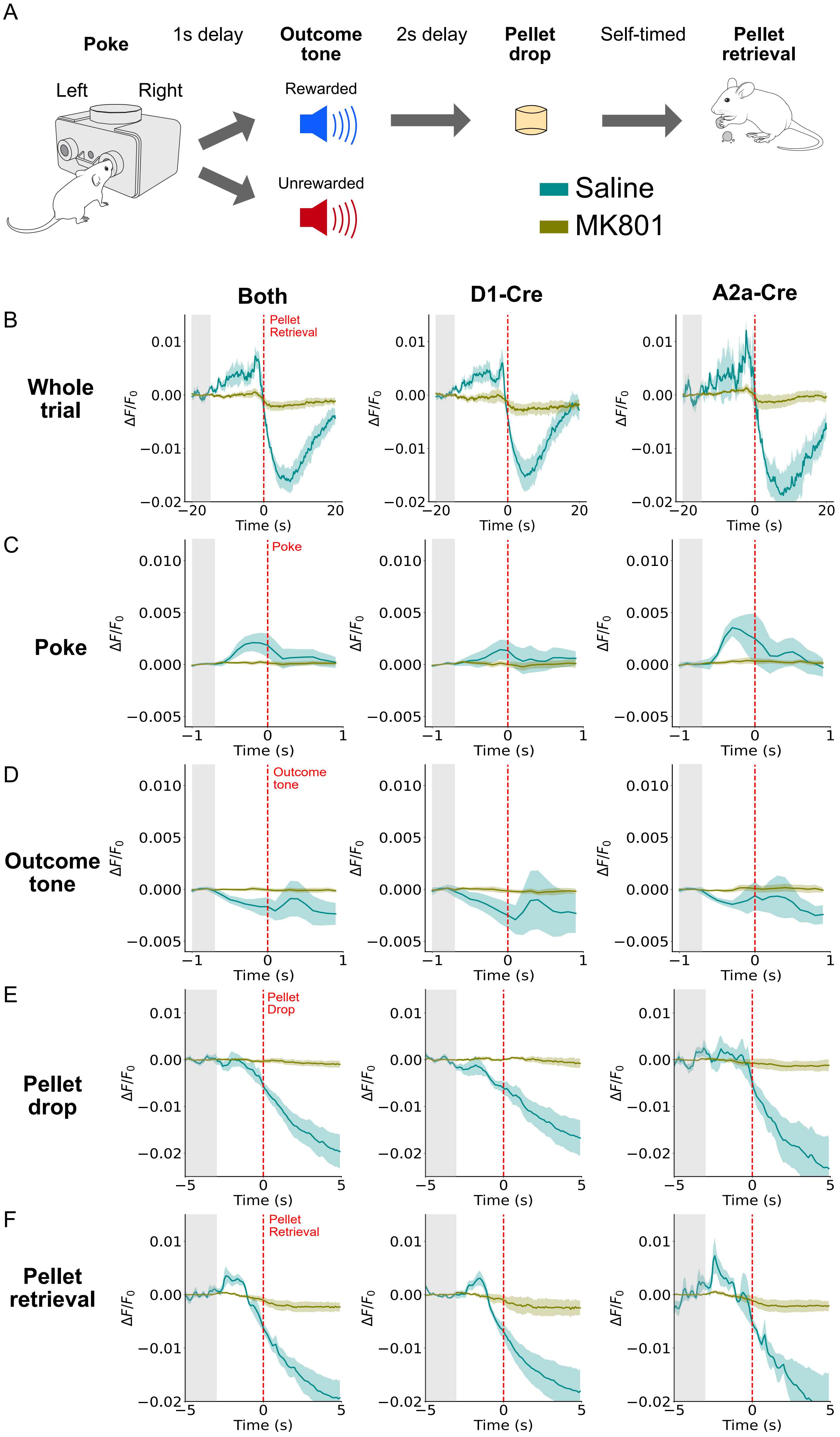


**Extended Data Figure 3. Systemic antagonism of NMDARs diminishes reward-driven behavioral calcium changes. (A)** Task structure. **(B-F)** Calcium activity in both MSN striatal populations (left, pooled data), D1-MSNs, or D2-MSNs during different stages of the trial, after saline or MK801 systemic injection. The gray bar indicates the baseline period to which each trace was normalized. **(B)** Entire trial, left panel identical to Figure 2H. **(C)** Aligned to poke, independent of side or outcome. **(D)** Aligned to outcome tone, independent of whether it was a rewarded or unrewarded trial. **(E)** Aligned to pellet drop. **(F)** Aligned to pellet retrieval.

### Extended Data Figure 4

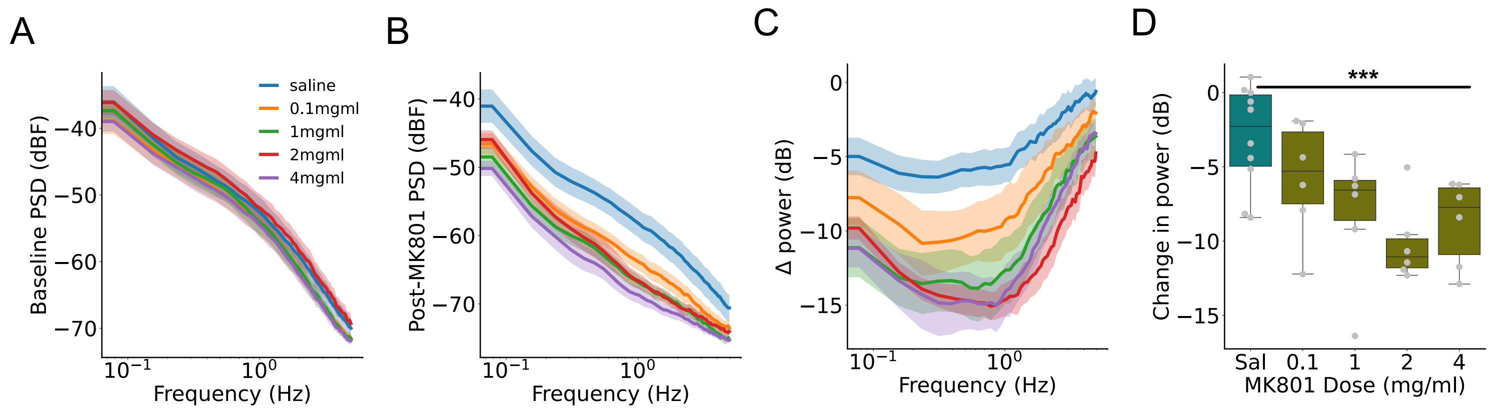


**Extended Data Figure 4. Antagonism of striatal NMDARs diminishes calcium signal power across frequencies in a dose-dependent manner. (A)** Baseline power spectrum density (PSD) prior to infusion of saline or MK-801. **(B)** PSD after infusion of MK-801. **(C)** Power difference between panels A and B. **(D)** Average change in power per mouse (from panel C).

### Extended Data Figure 5

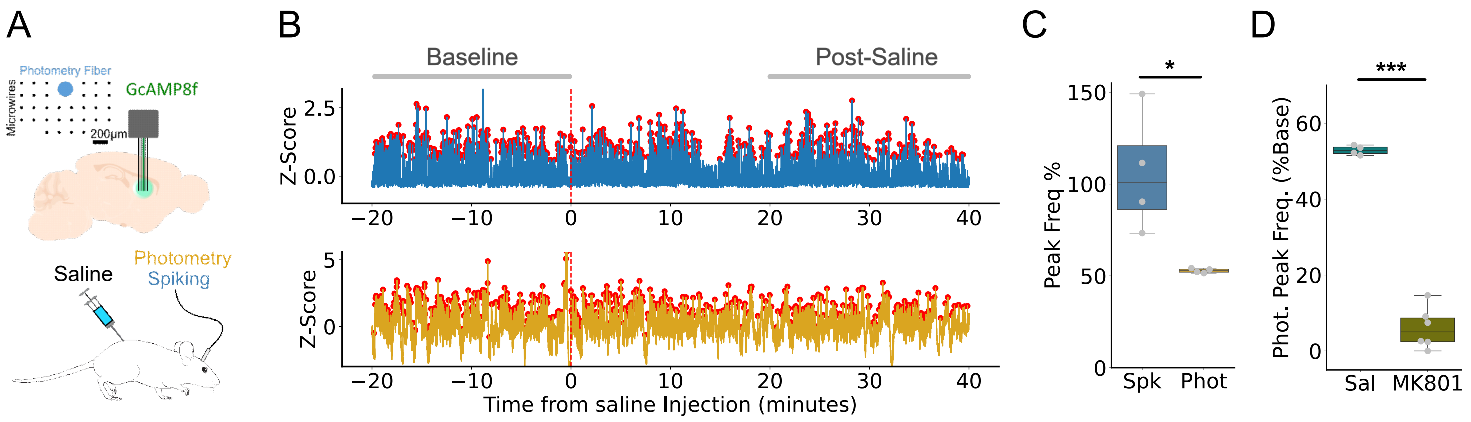


**Extended Data Figure 5. Calcium activity remains after systemic saline injection but not after MK801 injection. (A)** Simultaneous *in vivo* electrophysiology an calcium recordings before and after systemic saline injection. **(B)** Example trace of simultaneous average firing rate (top) and calcium activity (bottom) before and after saline injection. **(C)** Frequency of rapid increase in firing rate (Bursts) or calcium activity (Transient), normalized to the baseline period. **(D)** Calcium transient rate after saline or MK801 injection same calcium data as Panel (C), for saline injection, and Figure 3C, for MK801 injection.
